# Breakpoint models shown no evidence of thresholds in recreational response to increasing wildfire smoke in the American West

**DOI:** 10.1101/2022.04.21.489032

**Authors:** Matt Clark, Alexander Killion, Matthew A. Williamson, Vicken Hillis

## Abstract

Ambient wildfire smoke in the American West has worsened considerably in recent decades, while the number of individuals recreating outdoors has simultaneously surged. Wildfire smoke poses a serious risk to human health, especially during long periods of exposure and during exercise. Here we aggregate data on black carbon, a major component of wildfire smoke, and recreational visitation in 32 U.S. national parks from 1980 - 2019 to examine how visitors respond to wildfire smoke. We hypothesize that visitor response may exhibit a threshold effect where ambient smoke reduces visitation after a critical level, but not before. We develop a series of breakpoint models to test this hypothesis. Overall, these models show little to no effect of ambient smoke on visitation to the 32 parks tested, even when allowing for critical thresholds at the extreme upper ranges of the smoke data. This suggests that wildfire smoke does not significantly alter behavior of park attendance. This finding has implications for the management of recreation areas, public health, and climate change adaptation broadly.

## Introduction

Wildfires in the American West have increased dramatically in both frequency and size over the last 40 years [1,2]. Approximately half of this increase in burned area can be attributed to anthropogenic climate change [3]. Thus, wildfires in western forests are expected to worsen over the next 40 years as the climate continues to warm [2,4–9]. Accompanying the increase in wildfires in the American West, is a projected increase in wildfire smoke; fire-prone areas may experience a two-fold increase in wildfire smoke by 2100 [10,11].

Wildfire smoke is a complex mixture of gasses aerated by biomass combustion across the landscape [12]. While many of these compounds have been shown to cause health problems in humans, organic carbon and black carbon particulate matter less than 2.5 *μ*m in diameter (PM_2.5_) are particularly harmful to human health [13]. PM_2.5_ causes between 260,000 - 600,000 global deaths annually and significantly increases community mortality even under acute exposure, especially in areas experiencing outbreaks of COVID-19 infection [12,14–16]. While both PM_2.5_ and wildfire smoke generally are made up of organic material (i.e. organic carbon) and black carbon, black carbon is the most visible component [17]. Black carbon is particularly prominent in visible wildfire smoke, as it has high light absorption properties that contribute to the dark appearance of plumes [18].

In some regions of the American West, wildfires now account for 50% of all atmospheric PM_2.5_, compared to less than 20% in 2010 [19]. As any level of ambient PM_2.5_ increases community morbidity, public health agencies increasingly call for individuals to limit outdoor recreational behavior when wildfire smoke is present [20–22].

While calls to limit outdoor recreation during wildfire smoke events have become ubiquitous in the summertime American West, a survey of federal land managers indicated that they feel they have a shortage of information describing how air quality actually affects the recreational behavior of their visitors [23]. Reviews of scientific literature come to a similar conclusion, pointing to a dearth of research describing how individuals respond to low air quality (although see [24]) and how climate change is affecting the recreation landscape [25,26].

This article uses estimates of black carbon and visitation data collected from national parks in the American West from 1980-2019 to answer whether individuals alter their recreation behavior in response to ambient wildfire smoke. We hypothesize that the effect of ambient smoke on national park visitation may be nonlinear, only showing a measurable impact in the upper ranges of the observed smoke values. Threshold effects such as this are common in ecology and human behavioral sciences [27–29]. Toms and Lesperance (2003) show that breakpoint models are an effective tool to account for these thresholds and make inferences about the effect of a predictor before and after a particular threshold in natural systems [30]. Here we develop a series of three hierarchical breakpoint models to determine if a threshold effect is present in national park visitation response to ambient smoke and if so, what the effect of smoke is on visitation post-threshold.

## Materials and methods

### Data collection

#### Visitation data

We obtained monthly visitation data through the [National Park Service (NPS) Visitor Use Statistics Portal](https://irma.nps.gov/STATS/). These data are generally collected via car counters, permit information, and concessionaire reporting. Methods of collection vary from park to park. Unit specific information can be found on the National Park Service Visitor Use Statistics Portal. Although collection methods are inconsistent across the sample, all data was taken as reported. We retrieved monthly data for all national park units in the Intermountain and Pacific West regions for years 1980 through 2019.

While wildfires have seen a dramatic increase across the United States over the last four decades, changes in wildfire smoke have been most concentrated in the westernmost regions of the country [19]. In the contiguous United States, the western regions are the only areas with a significant smoke burden where the majority of the smoke they see originates from wildfires within their own borders [31]. Therefore, as to not confound our analysis with unpredictable smoke events which originate from distant wildfires, we limit our sample to the national parks in the American West (i.e. Intermountain and Pacific West NPS regions).

The 32 national parks included in the two regions in our sample have different high seasons for visitation. Wildfire smoke occurs year-round, but primarily in the Summer months. In some seasons, for some parks, there is a natural visitation drop driven by temperature alone that coincides with highest levels of wildfire smoke. To ensure that we estimate visitation changes driven by smoke rather than seasonality, we subset our data to only the three month high season associated with each park. For example, only Summer (June, July, Aug.) data were used for Yellowstone National Park and only Winter data (Dec., Jan., Feb.) were used for Death Valley National Park.

#### Smoke data

To detect times of high concentrations of black carbon associated with smoke, we used data from the second Modern-Era Retrospective analysis for Research and Applications [32]. MERRA-2 is a NASA atmospheric reanalysis that begins in 1980 and replaces the original MERRA reanalysis using an upgraded version of the Goddard Earth Observing System Model, Version 5 data assimilation system [33]. MERRA-2 provides mean monthly measurements of black carbon starting in 1980 at a spatial resolution of 0.625° x 0.5°. We used the monthly black carbon column mass density measurements (kg/m^−2^) and calculated the mean of those pixels that intersected national park boundaries to represent wildfire smoke at the park level; Fig 1 [34]. We also correlated the column black carbon values to the amount of black carbon only at the surface layer (i.e., the measurements closest to the ground; S1).

**Fig 1.**
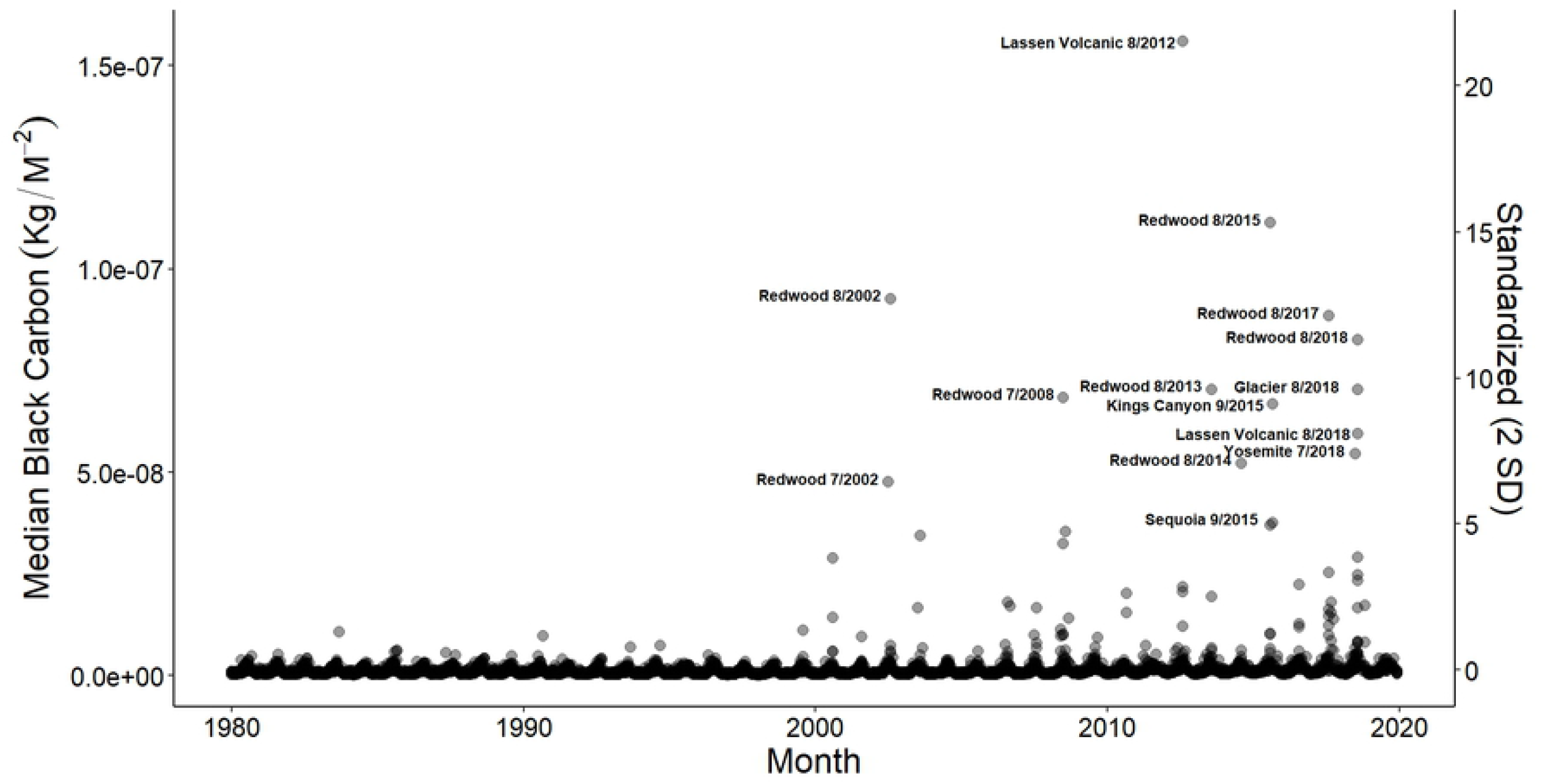
Smoke observations. Monthly smoke observations in each of the 32 national parks included in this study from 1980-2019. Monthly medians greater than 5 on our standardized scale (10 standard deviations above the mean) are labeled. The right-hand y-axis shows the standardized values of smoke referred to when describing the chosen critical thresholds. Smoke values on the left-hand y-axis are shown in kilograms per time-averaged two-dimensional meter.

### Analyses

Visitation to US national parks has increased sharply in recent years [35]. We first account for this underlying change in national park visitation from 1980-2019 by developing a hierarchical, temporal autoregressive model which we assess for within-sample predictive accuracy and build upon. This baseline autoregressive model Eq (1) predicts the visitation to a given park in a given month based on the recorded visitation in that park and month in the previous year. This formulation of an autoregressive model is commonly referred to as an AR(*k*) model, where *k* is the number of previous time periods used for prediction [36]. In this particular instance we chose *k* = 1 to reduce issues of collinearity and identifiability among autoregressive predictors [37].

We formulated this baseline autoregressive model using a Bayesian hierarchical framework. This approach accounts for between-park variation in visitation trends while acknowledging the interconnectedness of these trends through partial pooling [38]. For each park (*j*) we estimate a parameter value (***v***) representing the trend in visitation from one year to the next, in order to predict the visitation for each month in our data set (*i*). Finally, we estimate both global (*α*0) and park specific (***β*0**_*j*_) intercepts to yield equation 1 below. We modeled these data using a negative binomial distribution to allow us to estimate an overdispersion parameter (*ϕ*) rather than assuming the dispersion is equal to the mean. We fit this model using 7 Markov chains run for 5,000 iterations.

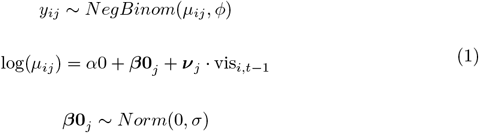

This baseline autoregressive model showed adequate diagnostic statistics, exhibiting ample mixing of Markov chains, 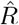 values equal to 1 for all parameter estimates, and the absence of divergent transitions after warm up. We examined the within-sample predictive capacity of this autoregressive-only model to be sure we sufficiently accounted for baseline changes in visitation before building on this model to test for the effects of smoke on visitation. This baseline model allowed us to account for approximately 37% of all variation in the data, with the bulk of the error occurring when actual visitation was very high or very low (Fig 2).

**Fig 2.**
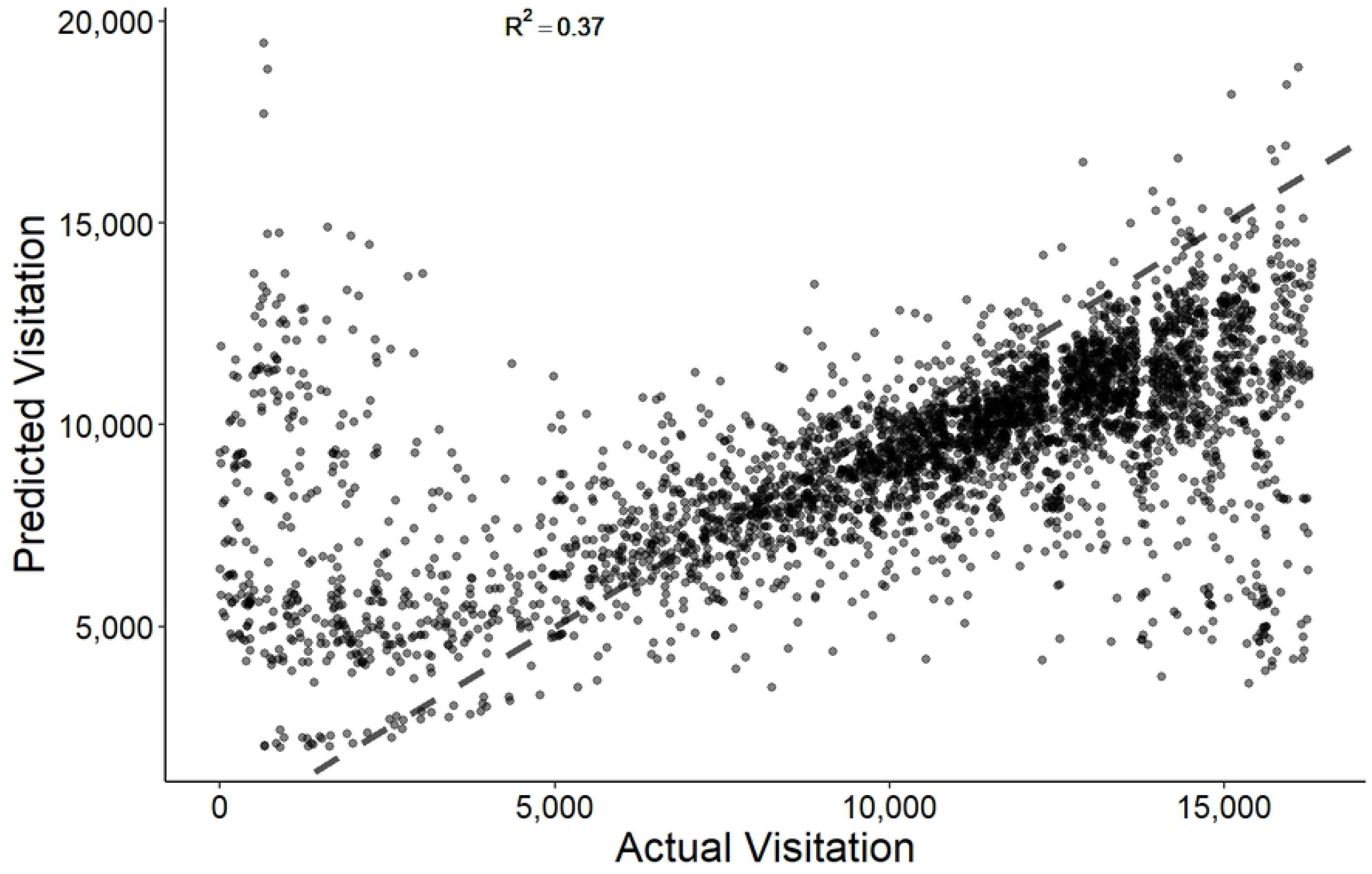
Autoregressive-only predictive accuracy. Scatterplot showing the within-sample predictive accuracy of the autoregressive baseline model used in this study. This model accounts for 37% of all variation in the data. Black dashed line shows an exact 1:1 relationship between the predicted visitation and actual visitation axes.

After confirming that the model above captures the baseline trends in national park visitation from 1980-2019, we built upon it to estimate if a critical threshold for visitation response to wildfire smoke exists in these data and if so, what the impact of wildfire smoke is on visitation post-threshold. To do this, we developed a series of three breakpoint models, allowing for three increasingly extreme breakpoints (i.e. critical thresholds). We standardized these smoke data, and thus the thresholds as well, by dividing by two standard deviations following the recommendation made by Gelman (2008) [39]. The three thresholds tested were 0.0, 0.5, and 1.0, which can thus be interpreted as the mean smoke value, one standard deviation above the mean, and two standard deviations above the mean respectively. These increasingly extreme thresholds contain the highest 45%, 8%, and 3% of observed smoke values out of the 3,744 data points in our sample. A visual comparison of the observed smoke values plotted on both the standardized and unstandardized scales can be seen in Fig 1.

We estimated the effect of smoke on visitation before (***β*1**) and after (***β*2**) each breakpoint (BP). Just as with the autoregressive term (***v***), we estimated unique parameter values for each park (*j*) in a hierarchical framework. The complete model used for each of the three breakpoint values is then as seen in Eq (2). Each model was run using 7 Markov chains for 5,000 iterations.

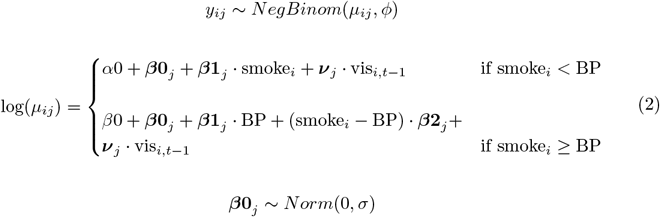

Following the recommendation given by Gelman et al. (2008) for producing stable, conservative estimates, we specified long-tailed regularizing priors for the autoregressive term, the global intercept value, and the overdispersion parameter [40]. We used normally distributed regularizing priors for estimating the standard deviation parameter (*σ*) and the pre and post-breakpoint smoke effects to allow for more efficient sampling while generating within-sample predictions at high smoke values.

As the goal of this paper is inference, we test just the suite of breakpoint models described above, which were developed *a priori* to evaluate our hypothesis that smoke would exhibit a non-linear threshold effect on park visitation [41]. We do not test our models against other candidate models using information criterion metrics or cross validation, as the objective of such tests is prediction rather than inference and we would not expect our models to be predictively valid strictly speaking [41,42].

## Results

All three breakpoint models showed adequate model diagnostics, including well-mixed Markov chains, 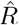 values of 1 for all estimated parameters, and the absence of divergent transitions after warm up [43]. We then take the coefficient estimates produced from our models as reliable for inference regarding our *a priori* hypothesis.

While some between-park variation exists, our overall study shows no evidence for threshold effects in recreational visitor response to wildfire smoke in national parks in the American West. Fig 3 and Fig 4 show the parameter estimates for the pre and post-breakpoint smoke effects respectively. In all three breakpoint models, only two parks have 90% credible intervals that do not overlap zero for slope 1, both of which have opposite signs (Fig 3). The post-breakpoint, slope 2, parameter estimates exclusively overlap zero at a 90% credibility interval (Fig 4). The uncertainty in these estimates increases dramatically as we use more extreme breakpoint values and the amount of data post-breakpoint decreases. We note however, that even parks with relatively certain estimates in the more extreme breakpoint models (0.5 & 1) still exclusively overlap 0, demonstrating no impact of ambient smoke on park visitation even at very dangerous levels.

**Fig 3.**
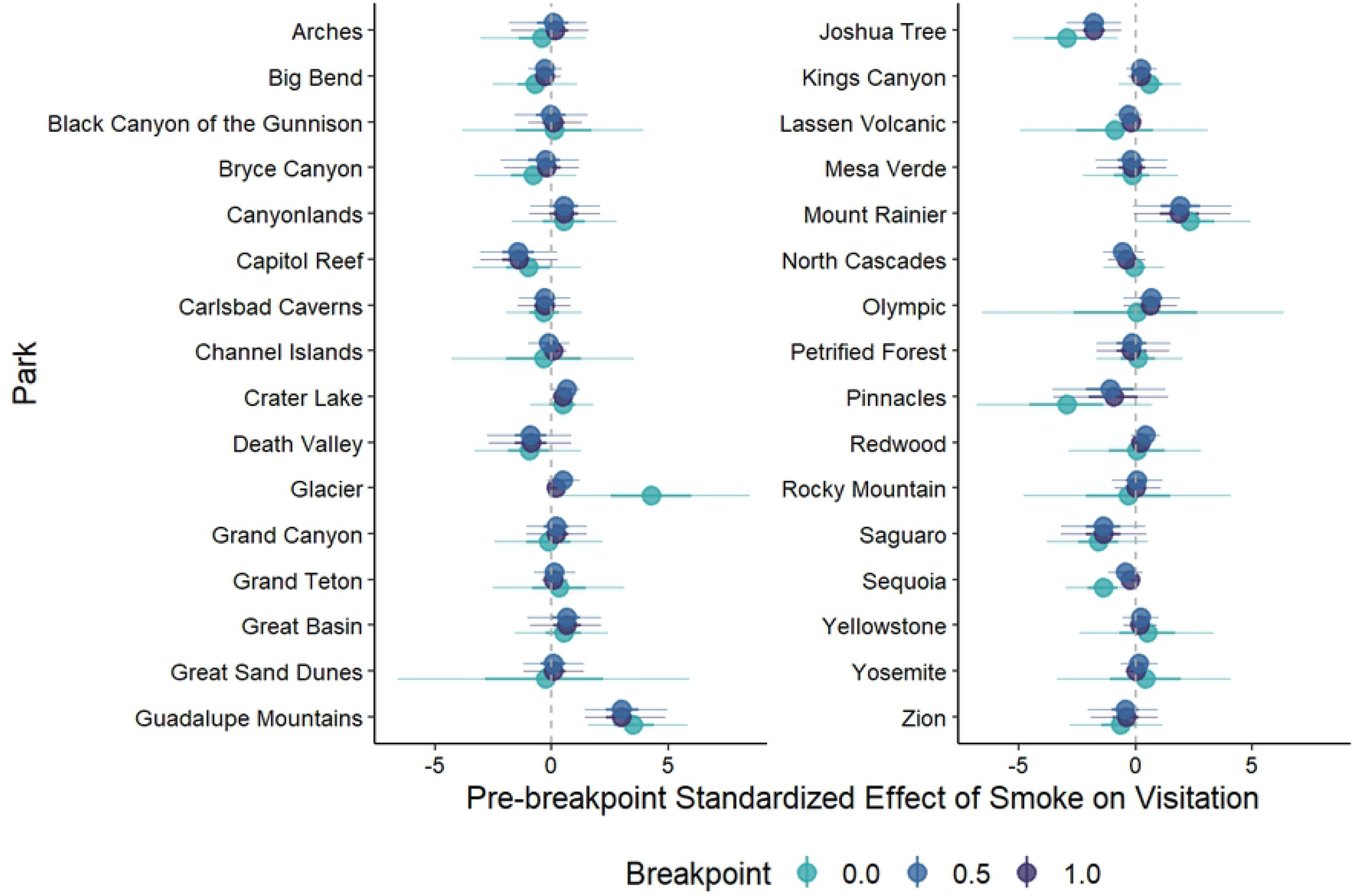
Slope 1 parameter estimates. Posterior parameter estimates for slope 1 for each of the 32 national parks included in this study for each of the three breakpoint models. Points represent the median prediction. Thick lines show the 50% credibility intervals and thin lines show the 90% intervals.

**Fig 4.**
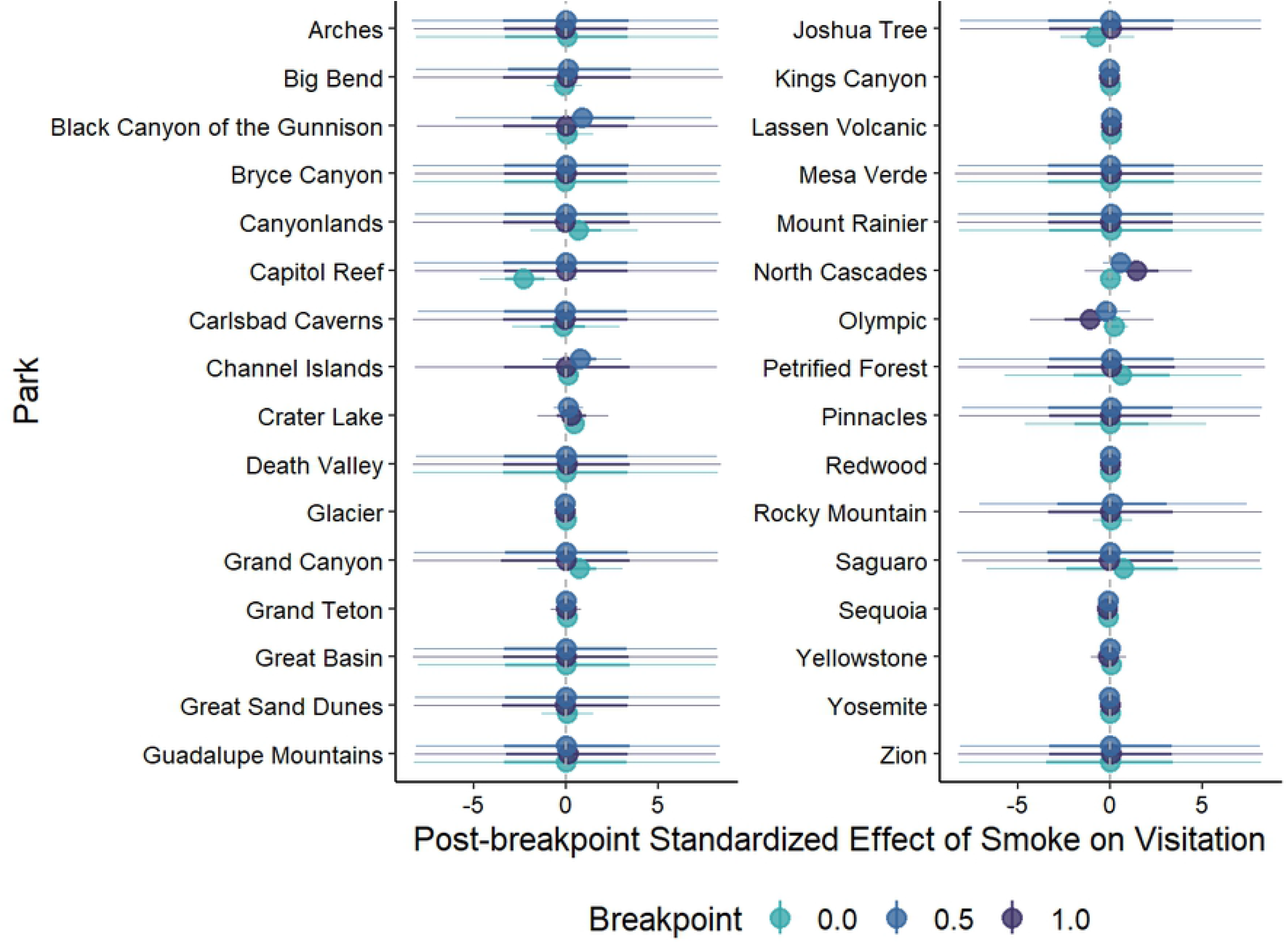
Slope 2 parameter estimates. Posterior parameter estimates for slope 2 for each of the 32 national parks included in this study for each of the three breakpoint models. Points represent the median prediction. Thick lines show the 50% credibility intervals and thin lines show the 90% intervals.

Visually examining the marginal effect of ambient smoke on park recreational visits provides a similar intuition as above (Fig 5). We see little difference in the effect of smoke before and after the hypothesized thresholds. In addition, we descriptively show high levels of visitation even under very high levels of ambient smoke. In Redwood National Park for example, we see visitation within the normal range even during the August 2018 wildfire events, which produced smoke levels over 30 times the total sample standard deviation.

**Fig 5.**
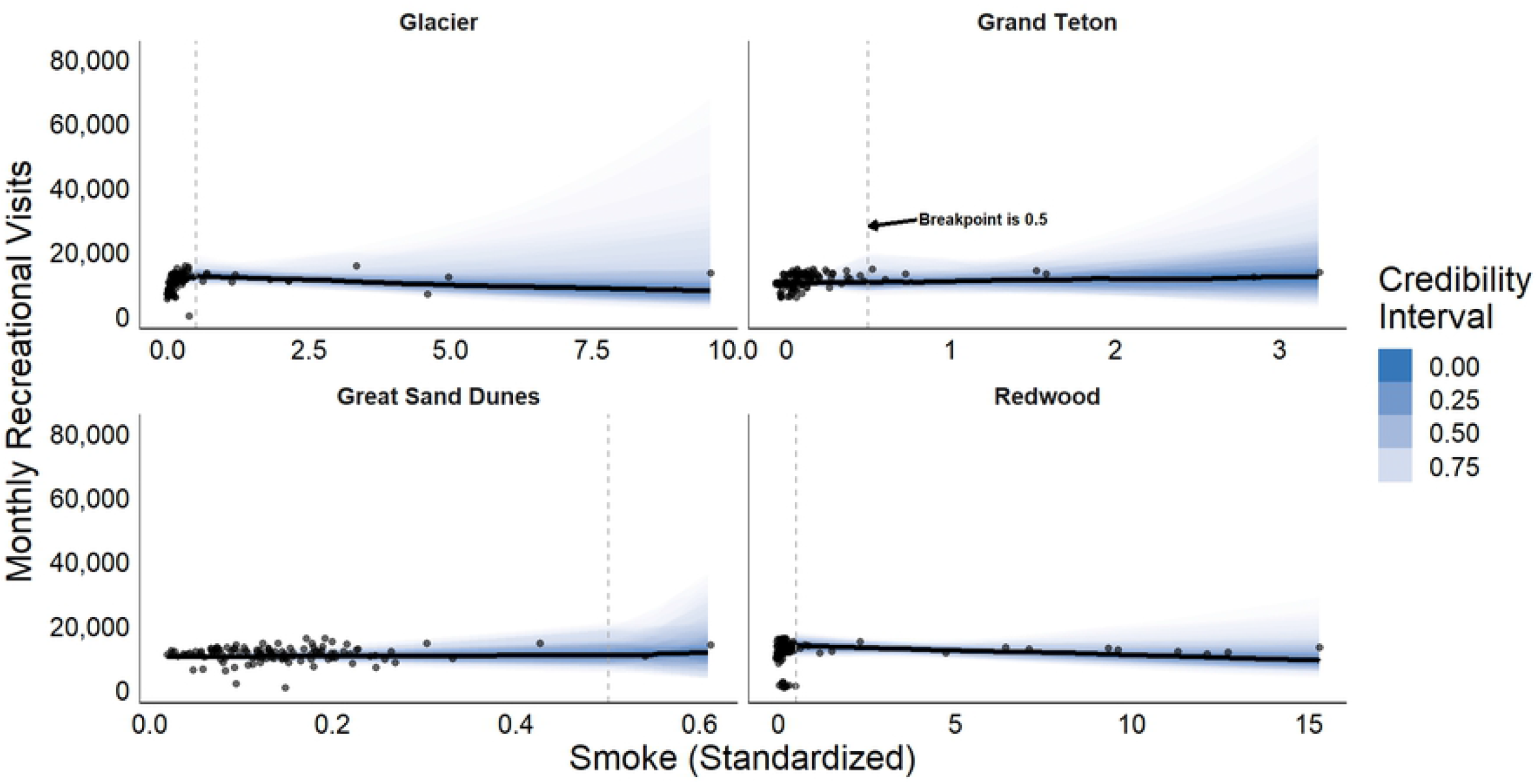
Marginal effects of slope on visitation. Marginal effect of ambient smoke on monthly park visitation for four of our study parks with the highest median smoke values from 1980 to 2019. Parks displayed are Glacier National Park, Grand Teton National Park, Great Sand Dunes National Park & Preserve, and Redwood National Park. For simplicity, we show just the model with the middle breakpoint value (0.5 on our standardized scale). Blue shading represents the full range of posterior predictions for visitation given the mean value for each autoregressive term in the model. Intensity of shading shows the credibility interval for each prediction, with the middle black line showing the median prediction at each value of smoke.

Given the clear lack of threshold effects in visitation response to ambient wildfire smoke, we conducted an exploratory analysis on the overall effect of smoke on visitation in our study parks (i.e. without a breakpoint). We formulate this exactly as in the pre-breakpoint slope in equation 2. We again ran this model for 5,000 iterations using 7 Markov chains, which exhibited adequate mixing, a lack of divergent transitions after warmup, and 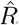 values of 1 for all estimated parameters.

The parameter estimates for the overall effect of ambient smoke on visitation response in all 32 parks in our sample are considerably more narrow than the estimates produced via the breakpoint models (Fig. 6). Still, the 90% credibility intervals estimating the smoke effect for each park exclusively overlap 0, indicating no overall effect, further supporting our findings above. Intuitively, the within-sample *R*^2^ remains equal to 0.37 after accounting for the overall smoke effects, showing no improvement compared to the autoregressive only model. This suggests that ambient smoke in national parks in the American West is not driving any measurable change in visitation even at sustained and dangerous levels. We thus conclude that visitors are not altering recreation behavior in response to smoke and therefore no overall effect or critical threshold for adaptation is detectable in these data.

**Fig 6.**
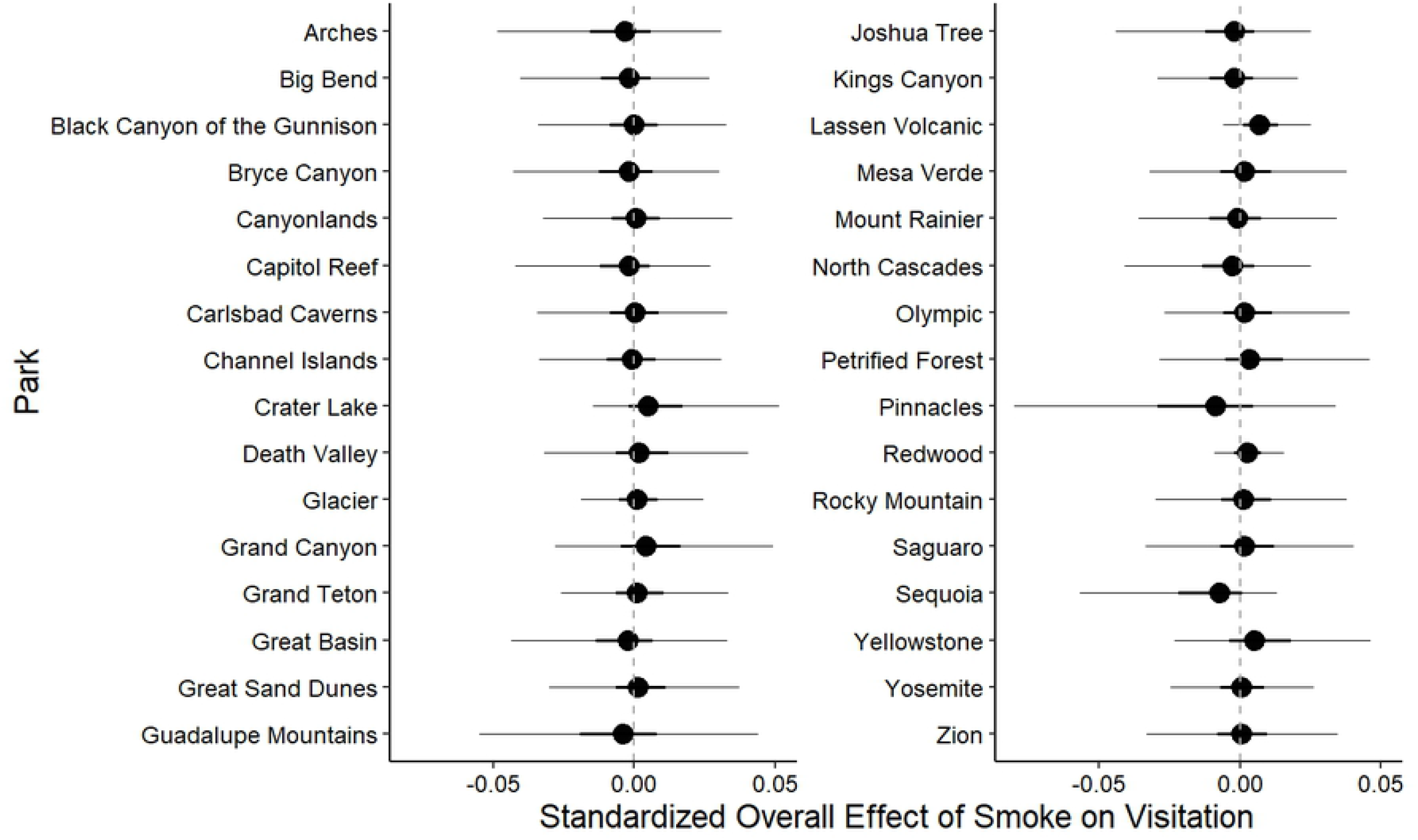
Overall model parameter estimates. Posterior parameter estimates for the overall effect of smoke on visitation for each of the 32 national parks included in this study. Points represent the median prediction. Thick lines show the 50% credibility intervals and thin lines show the 90% intervals.

## Discussion

We did not detect visitor adaptation to increasing wildfire smoke in national parks in the American West. This result is troubling both specifically for visitor health in U.S. national parks and for climate change adaptation broadly. As discussed above, wildfire smoke significantly increases community morbidity even under acute exposure. In concert with showing no overall trends in behavioral adaptation, these data provide specific instances of historic smoke events where visitation did not deviate from normal (Fig 5).

The highly variable nature of wildfires, their relationship to climate change, and the great distances that smoke can travel during and after wildfire events make it difficult for individuals to plan around smoke events [31,44,45]. In conjunction, U.S. national parks draw a great number of non-local visitors, many of whom are coming from other states or countries and are likely visiting for the first time [46]. We speculate that these visitors are less likely to change their plans due to wildfire smoke than individuals recreating locally or repeatedly in one location. We propose that these unique features of national parks and wildfire smoke make visitors particularly unable to adapt to changing climatic conditions.

Based on our findings here, we suggest that a regional or national level policy limiting visitation during dangerous smoke events may be necessary to protect would-be visitors to U.S. national parks [47]. Presently, there is considerable variation in the way states react and plan for climate change and associated hazards [48].

Considering our findings more generally, humans must adapt quickly to the new realities of our increasingly variable climate if we are to continue to thrive over the coming decades [49]. Climate change is already dramatically altering the social-ecological landscapes in which humans have learned to operate [50]. This research contributes to a broader body of work showing that despite increasing awareness, as a society we still fail to respond adequately [51]. Even when adaptive strategies and their benefits are known, we do not translate strategies into action [52]. While research has shown that extreme events tend to spur climate adaptation on the part of governments, individual response is much less consistent [53–55]. Therefore, while we speculate that the system of recreational visitation to U.S. national parks may be particularly problematic, we suppose that the trends observed in this study may be characteristic of individual response to many changing environmental conditions.

### Limitations and future research

A key limitation of this study is the lack of fine scale visitation data to US national parks. Daily visitation data would allow us to identify a more complete picture of if and how visitors respond to smoke events day to day. Current data at the monthly scale allows us to investigate the big picture trends in visitor behavior, but future research would benefit greatly from daily count data. It is possible that individuals postpone visits to national parks by days or weeks when smoke is present, behavior which our current study would be unable to capture.

It is also possible that this trend would not be observed for recreation on all public lands or for all climate change-induced hazards. Other paths for future research may therefore be to investigate if the trend shown here holds for other public lands with different visitor profiles and for other hazards.

## Conclusion

The results presented here indicate that individuals are not modifying their behavior to adapt to worsening wildfire smoke events. It is unclear if this finding is representative of individual climate adaptation generally or is unique to this system. Regardless of the generality of our findings, this study has specific implications for management of U.S. national parks. Future research may confirm or negate the trend shown here in individual climate adaptation broadly. This will have considerable implications for the future of all communities living with climate-induced hazards.

## Supporting information

**S1.**
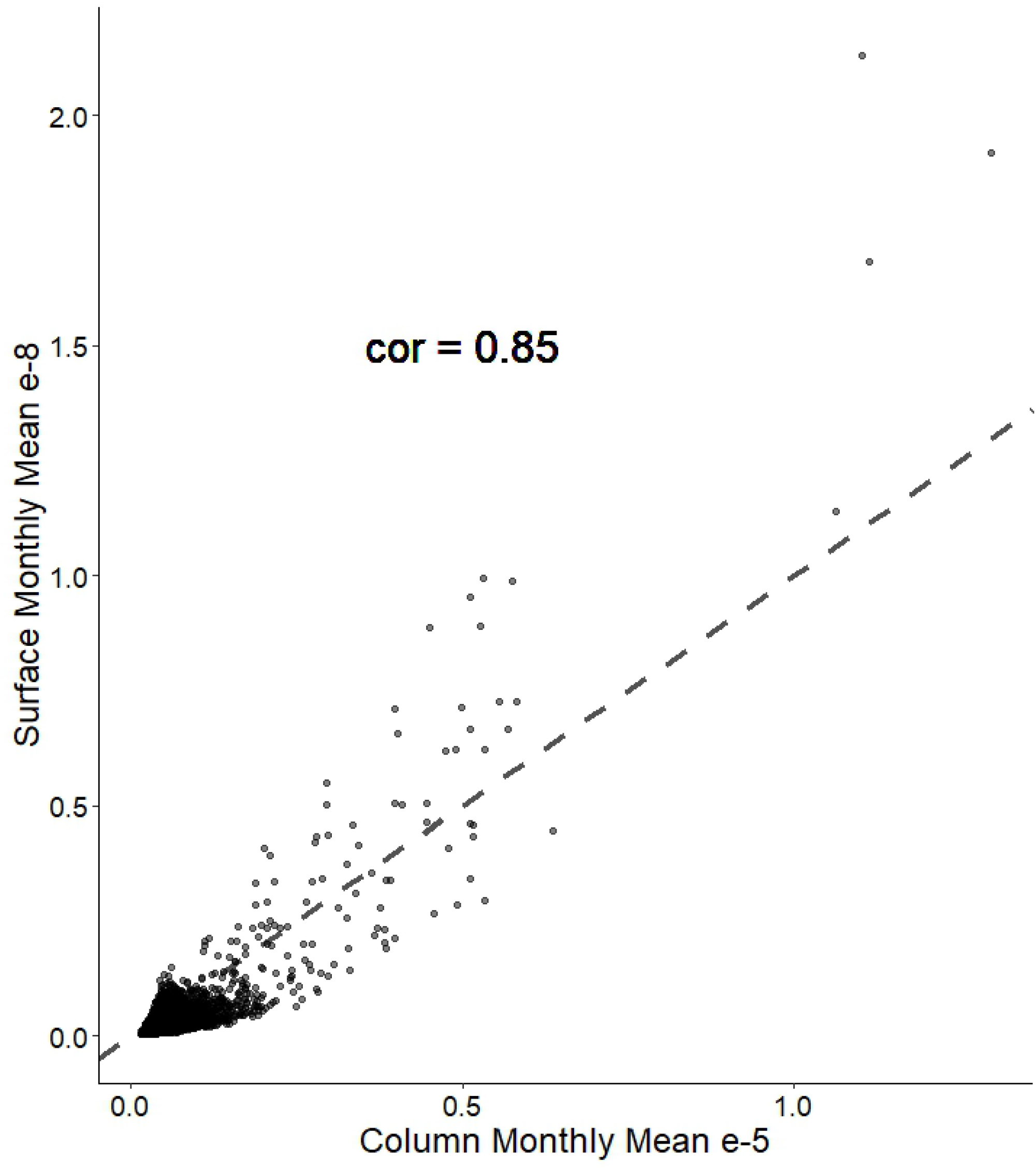
Correlation between surface level and column mean black carbon measurements. Scatterplot showing the correlation between measurements of the surface level and the column-wide black carbon densities. Densities are displayed in (kg/m^−2^). We use just surface level density measurements in our analyses. The surface and column-wide densities are correlated at a value of 0.85.

## Acknowledgments

We would like to thank Dr. Mojtaba (Moji) Sadegh for initial thoughts on the selecting appropriate smoke data for this analysis. Additionally, we would like to thank the entire ”EcoStats” group at Boise State University for ongoing support on all things related to quantitative ecology.

